# ZEB2 signaling is essential for ureteral smooth muscle cell differentiation and maintenance

**DOI:** 10.1101/2025.02.23.639741

**Authors:** Sudhir Kumar, Xueping Fan, Harshita Pattam, Kun Yan, Easton Jinhun Liaw, Jiayi Ji, Emily Zaltz, Paul Song, Yuqiao Jiang, Yuriko Nishizaki, Yujiro Higashi, Chen-Leng Cai, Weining Lu

## Abstract

Mowat-Wilson Syndrome (MWS) is a multiple congenital anomaly syndrome caused by mutations in the *ZEB2,* which plays a critical role in cell fate determination and differentiation during development. Congenital anomalies of the kidney and urinary tract (CAKUT) have been reported in MWS patients. However, the role of ZEB2 in urinary tract development and the cellular and molecular mechanism underlining the CAKUT phenotypes in MWS remains unknown. We performed ZEB2 protein expression analysis in the developing mouse ureter. We generated *Zeb2* ureteral mesenchyme-specific conditional knockout mice by crossing *Zeb2* floxed mice with Tbx18Cre mice (*Zeb2* cKO) and analyzed the urinary tract phenotypes in *Zeb2* cKO mice and wild-type littermate controls by gross and histological examination. Ureteral cellular and molecular phenotypes were studied using TAGLN, ACTA2, FOXD1, POSTN, CDH1, TBX18, and SOX9 ureteral cell-specific markers. We found that ZEB2 is expressed in TBX18^+^ ureteral mesenchymal cells during mouse ureter development. Deletion of *Zeb2* in developing ureteral mesenchymal cells causes hydroureter and hydronephrosis phenotypes, leading to obstructive uropathy, kidney failure, and early mortality. Cellular and molecular marker analyses showed that the TAGLN^+^ACTA2^+^ ureteral smooth muscle cells (SMCs) layer is not formed in *Zeb2* cKO mice at E15.5, but the FOXD1^+^ and POSTN^+^ tunica adventitia cells layer is significantly expanded compared to wild-type controls. CDH1^+^ urothelium cells are reduced considerably in the *Zeb2* cKO ureters at E15.5. Mechanistically, we found that *Zeb2* cKO mice have significantly decreased TBX18 expression but an increased SOX9 expression in the developing ureter at E14.5 and E15.5 compared to wild-type littermate controls. Our results show that ZEB2 is essential for ureter development by maintaining ureteral mesenchymal cell differentiation into normal ureteral SMCs. Our study also shed new light on the pathological mechanism underlying the developmental abnormalities of the urinary tract phenotypes in MWS patients.

**Author Summary:** Hydroureter and hydronephrosis are common congenital anomalies with high incidence (1:100 to 1:500) that can lead to obstructive uropathy and renal failure in the pediatric population. Mowat-Wilson Syndrome (MWS), caused by heterozygous mutations in the *ZEB2* gene, is a genetic disease with multiple congenital developmental defects, including hydroureter and hydronephrosis. However, the molecular function of ZEB2 in ureter development and pathogenesis of hydroureter and hydronephrosis remains unknown. Here, we show that ZEB2 is expressed in the developing ureteral mesenchymal cells, and deletion of ZEB2 in ureteral mesenchymal cells leads to ureteral smooth muscle cell loss, which is replaced by tunica adventitia cells causing hydroureter and hydronephrosis phenotype. Hence, our work not only demonstrates the critical role of ZEB2 in ureter development but also provides molecular mechanisms of urinary tract anomalies in MWS patients.

## Introduction

The mammalian ureters consist of three primary cell layers: the outer layer of fibrocytes of the tunica adventitia and lamina propria, the middle smooth muscle cell (SMC) layer, and the inner urothelial layer [1–4]. Between E14.5 and E16.5, the TBX18^+^ ureteric mesenchymal cells differentiate into the middle SMC layer and form a fully functional ureter at E16.5 before urine production from the kidney [5]. Any abnormality in ureteral SMC development impairs the formation of a functional ureter, leading to ureter dilation, hydroureter, hydronephrosis, and obstructive uropathy [1–3]. Hydroureter and hydronephrosis have a high incidence (1:100 to 1:500) in the pediatric population and are a leading cause of obstructive uropathy and renal failure [6, 7].

Mowat-Wilson Syndrome (OMIM #235730; MWS), caused by heterozygous mutations or deletions in the *ZEB2* gene, is a multiple congenital anomaly syndrome characterized by atypical face, moderate-to-severe mental retardation, epilepsy, and variable congenital malformations including Hirschsprung disease (HSCR), congenital heart defects, agenesis of the corpus callosum, and eye anomalies [8–11]. Over 50% of MWS patients have urogenital/renal anomalies, including duplex kidney, pelvic kidney, vesicoureteral reflux (VUR), hydroureter, and hydronephrosis [12–14]. Although many MWS patients develop hydroureter and hydronephrosis, the molecular function of ZEB2 in ureter development and the pathogenesis of CAKUT has not been studied.

ZEB2 is a zinc-finger E-box–binding (ZEB) protein family member. ZEB2 plays a critical role in cell fate determination and differentiation of embryonic stem cells, including hematopoietic progenitors and neural progenitor cells [15–22]. Previously, we demonstrated that loss of *Zeb2* in nephron progenitors leads to glomerulocystic kidney and renal failure [23]. We also found that ZEB2 controls FOXD1^+^ stromal progenitor cell differentiation, and loss of ZEB2 in kidney stroma progenitors leads to kidney fibrosis [24]. In this study, we hypothesize that ZEB2 also plays a critical role in ureter development, and the loss of ZEB2 in the developing ureter leads to abnormal ureter morphogenesis and hydronephrosis. To test this hypothesis, we analyzed the ZEB2 expression in developing mouse ureter and generated ureter-specific *Zeb2* conditional knockout mice to uncover its role in ureter development.

Our results show that ZEB2 is highly expressed in the TBX18^+^ ureteral mesenchymal cells. Deletion of *Zeb2* in mouse ureteral mesenchymal cells using *Tbx18Cre*^+^ led to hydroureter, hydronephrosis, and early mortality. Cellular marker analyses showed that ureteral mesenchyme-specific *Zeb2* conditional knockout mice (also called *Zeb2* cKO in this paper) had no TAGLN^+^ACTA2^+^ ureteral smooth muscle cells (SMCs) but instead, had an expanded layer of FOXD1^+^POSTN^+^ ureteral tunica adventitia cells during early ureter development. Mechanistically, we found that *Zeb2* knockout ureters had a reduced TBX18 expression with an upregulated SOX9 expression, which are critical transcriptional factors for ureteral SMC development. Our results indicate that ZEB2 is essential for the differentiation of ureteral SMCs, and loss of ZEB2 leads to depletion in the ureteral SMC layer that is replaced by tunica adventitia layer, eventually leading to hydroureter, hydronephrosis, obstructive uropathy, and renal failure.

## Result

### ZEB2 is expressed in TBX18^+^ ureteral mesenchymal cells in the developing ureter

To elucidate the role of *Zeb2* during ureter development, we first checked the expression of ZEB2 using immunostaining and found that ZEB2 is highly expressed in the mouse-developing ureters (Figure 1A). ZEB2 expression in the ureter was also confirmed by analyzing a ZEB2-EGFP knock-in reporter mouse that was previously generated [25] (Figure 1A). To determine the cell-specific expression of the ZEB2, we performed marker analysis by double immunostaining of ZEB2 with ureteral mesenchymal cell marker TBX18 and urothelial cell marker cadherin-1 (CDH1). The results revealed that ZEB2 was co-localized with TBX18 but not with CDH1 (Figure 1B-C), suggesting that ZEB2 is expressed by TBX18^+^ ureteral mesenchymal cells during the ureter development.

**Figure 1.**
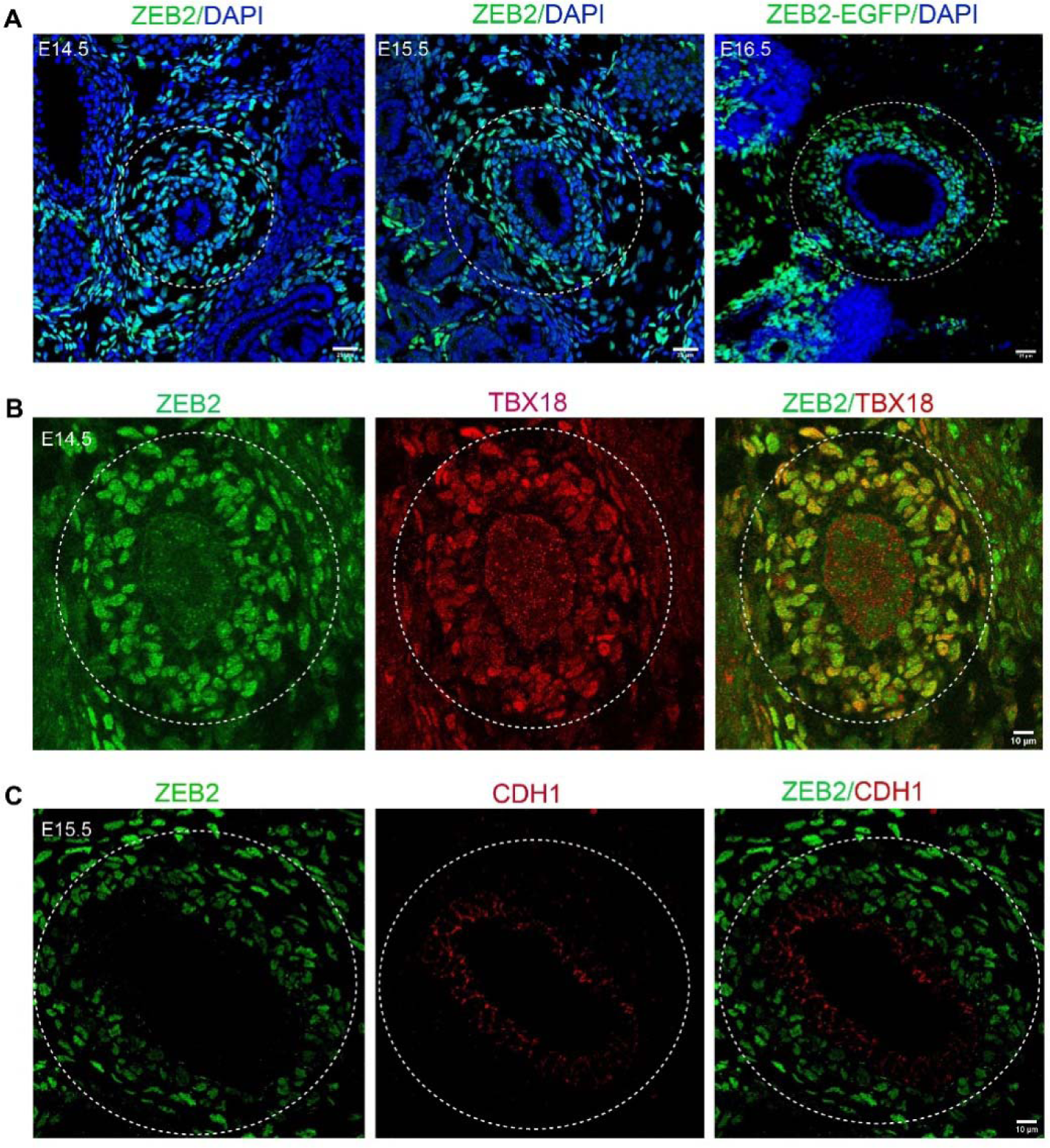
ZEB2 is expressed in TBX18^+^ ureteral mesenchymal cells in the developing ureter. (A) The left two panels show ZEB2 (green) expression at two developing stages (E14.5-E15.5) in the mouse ureter. The right panel shows a knock-in ZEB2-EGFP (green) expression. (B) ZEB2 (green) is co-localized with ureteral mesenchymal progenitor cell marker TBX18 (red) in the mouse developing ureter at E14.5. (C) ZEB2 (green) is not co-localized with CDH1 (urothelial cell marker) in the mouse developing ureter. Dotted lines encircle the ureter. Scale bars: 25 µm (A) and 10 µm (B and C).

### Loss of *Zeb2* in TBX18^+^ ureteral mesenchymal cells leads to hydroureter, hydronephrosis, and early mortality

To understand the function of ZEB2 in TBX18^+^ ureteral mesenchymal cells, we generated *Zeb2* conditional knockout mice, *Zeb2*^flox/flox^;*Tbx18Cre*^+^ (*Zeb2* cKO), by crossing *Zeb2* floxed mice [26] with *Tbx18Cre^+^*mice [27]. The deletion of *Zeb2* in ureteral mesenchymal cells was confirmed by immunostaining, which showed no ZEB2 signals in the *Zeb2* cKO mouse ureter compared to wild-type controls (Figure 2A). *Zeb2* cKO mice were born in the Mendelian ratio (Table 1), but most homozygous *Zeb2* cKO mice died within a few hours after birth and only a few survived up to 3 weeks of age (Table 1). The 3-week-old *Zeb2* cKO mice were runted and hunched (Figure 2B) and developed hydroureter and hydronephrosis phenotypes (Figure 2C-D). We also analyzed newborn mice (P0) and found that homozygous *Zeb2* cKO newborn pups also developed hydroureter and hydronephrosis phenotypes as well (Figure 2E-G). Our data suggest that in the absence of ZEB2, ureters develop obstructive uropathy, which eventually leads to kidney failure and early mortality.

**Figure 2:**
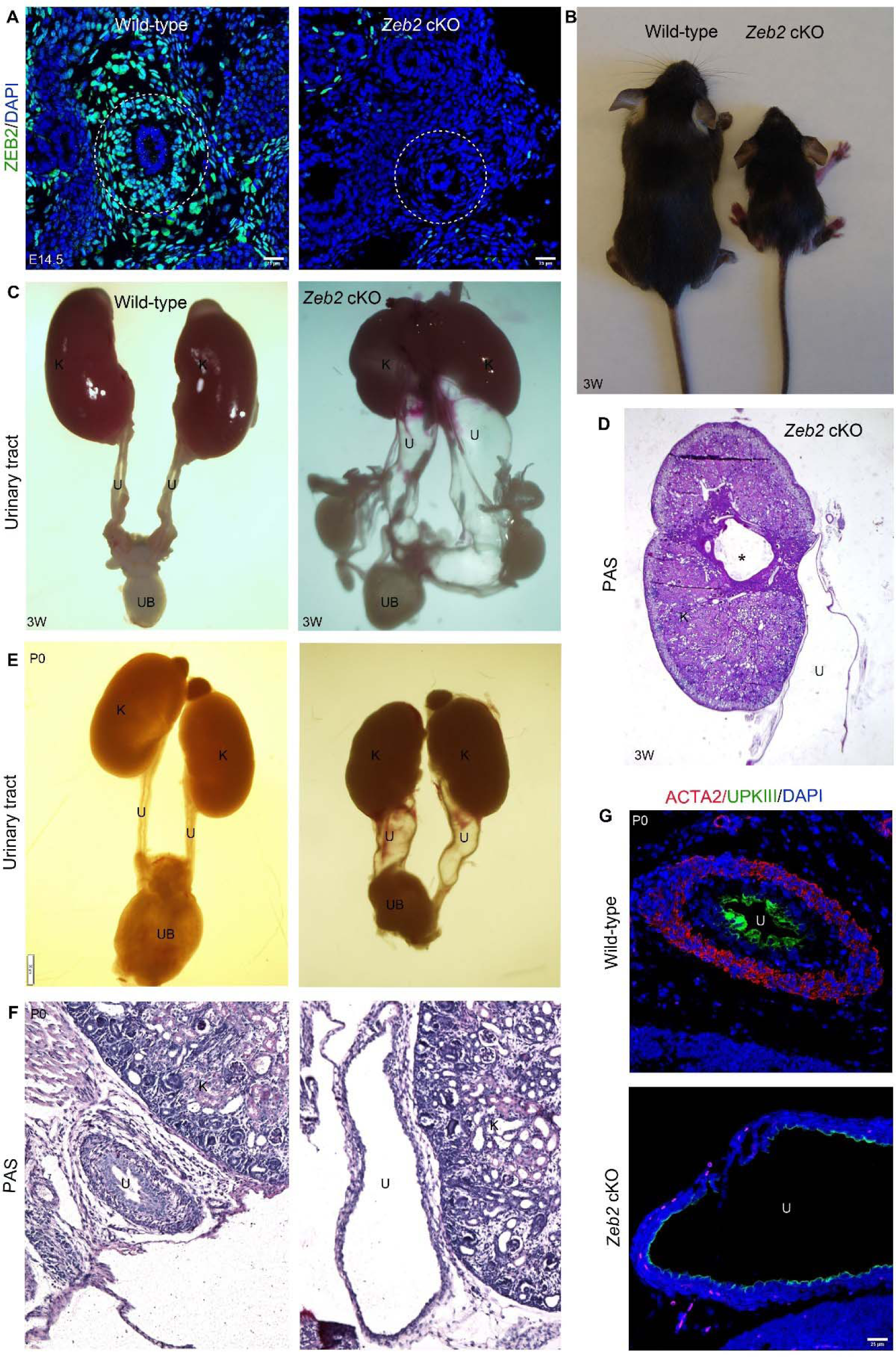
Loss of *Zeb2* in TBX18^+^ ureteral mesenchymal cells leads to hydroureter, hydronephrosis, and early mortality. (A) ZEB2 (green) expression is absent in *Zeb2*^flox/flox^;*Tbx18Cre*^+^ knockout mice (*Zeb2* cKO) as compared to wild-type littermate. (B) 3-week-old *Zeb2* cKO mice were runted with smaller body sizes as compared to wild-type control littermates. (C) 3-week-old *Zeb2* cKO mice developed hydroureter and hydronephrosis. (D) PAS staining of the kidney of a 3-week-old *Zeb2* cKO mouse showing dysplastic kidney with dilated ureter and hydronephrosis (*). (E) Newborn (P0) *Zeb2* cKO mice develop hydroureter and hydronephrosis. (F) Histology of a newborn (P0) *Zeb2* cKO mouse ureter with hydroureter as compared to wild-type littermate controls. (G) Immunofluorescence staining of ACTA2 (smooth muscle cell marker) and UPKIII (urothelium marker) shows abnormal ureter development with significant ureter dilation in the newborn (P0) *Zeb2* cKO mouse as compared to the wild-type littermate controls. u: ureter, k: kidney, UB: urinary bladder. Dotted lines encircle the ureter. Scale bars: 25 µm (A and G); 100 µm (D and F).

**Table 1:** *Zeb2* cKO mice were born in a Mendelian ratio but died after birth.

| Genotype | <u>3 weeks (*n* = 79)</u> |  | <u>P0 (*n* = 78)</u> |  |
| --- | --- | --- | --- | --- |
|  | <u>Observed</u> | <u>Expected</u> | <u>Observed</u> | <u>Expected</u> |
| *Zeb2*<sup>flox/flox</sup>;*Tbx18Cre*<sup>+</sup> (*Zeb2* cKO) | 4 (5%) | 25% | 20 (26%) | 25% |
| *Zeb2*<sup>flox/+</sup>;*Tbx18Cre*<sup>+</sup> (*Zeb2* het) | 28 (35%) | 25% | 22 (28%) | 25% |
| *Zeb2*<sup>flox/flox</sup>;*Tbx18Cre*<sup>-</sup> (WT) | 23 (29%) | 25% | 18 (23%) | 25% |
| *Zeb2*<sup>Flox/+</sup>;*Tbx18Cre*<sup>-</sup> (WT) | 24 (31%) | 25% | 18 (23%) | 25% |

### Ureteral mesenchymal progenitors fail to differentiate into smooth muscle cells in the absence of ZEB2

During the ureter development, TBX18^+^ ureteral mesenchymal cells differentiate into smooth muscle cells (SMC), important for functional ureter formation and prevents ureter dilation after urine production [2, 5]. To check the SMC formation in the developing ureter of *Zeb2* cKO mice, we examined two established SMC markers, the actin alpha 2, smooth muscle (ACTA2, Gene ID: 59), and transgelin (TAGLN, Gene ID: 6876) [4, 28]. Pan-cytokeratin was used to demarcate the urothelial cells [29]. We found that wild-type mice formed a normal ACTA2^+^TAGLN^+^ SMC layer at E15.5, however, this layer was absent in the age-matched *Zeb2* cKO mice (Figure 3A-B). TBX18^+^ ureteral mesenchymal cells also give rise to tunica adventitia layer that expressed marker genes forkhead box D1 (FOXD1, Gene ID: 2297) and periostin (POSTN, Gene ID: 10631) [4]. We, therefore, asked if there is any change in tunica adventitia ureter layer development in the absence of ZEB2. We examined the tunica adventitia cells by specific antibody immunostaining and found that there was an expanded FOXD1^+^POSTN^+^ tunica adventitia layer in the developing ureter of *Zeb2* cKO mice as compared to their wild-type littermate controls (Figure 3C-D). Urothelial cell differentiation was also reduced in *Zeb2* cKO E15.5 developing ureter as evidenced by a significantly reduced CDH1-expressing developing urothelial cells (Figure 4A-B) and pan-cytokeratin staining (Figure 3A-B). This is likely due to abnormal ureteral mesenchymal differentiation as the ureteral mesenchymal cells crosstalk with urothelial cells during ureter development [30]. Taken together, our data suggest that ZEB2 is essential for the normal differentiation of the ureteral mesenchymal cells into SMCs to form functional ureters, and loss of ZEB2 causes the absence of SMCs in developing ureter leading to hydroureter and hydronephrosis phenotypes.

**Figure 3.**
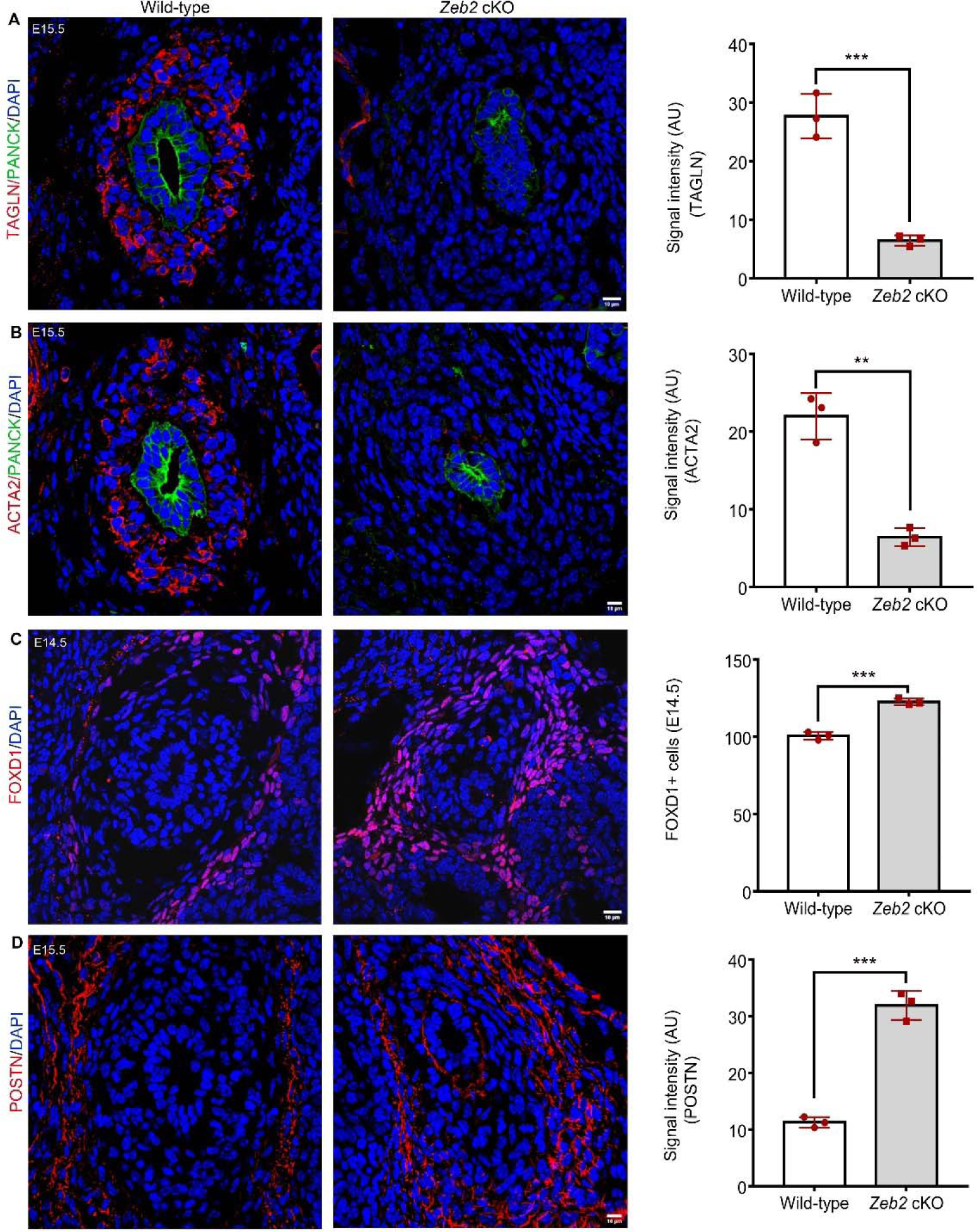
Ureteral mesenchymal progenitors fail to differentiate into smooth muscle cells in the absence of ZEB2. (A, B) Ureteral smooth muscle cells are formed in the wild-type mice at E15.5 specified by smooth muscle markers TAGLN (A) and ACTA2 (B). There are no TAGLN^+^ and ACTA2^+^ ureteral smooth muscle cells in *Zeb2* cKO mice. Pan-cytokeratin (Pan-cyto) is used to mark urothelial cells. (C, D) Compared to wild-type littermate controls, ureteral tunica adventitia cell layers labeled by POSTN+ and FOXD1+ markers are expanded in Zeb2 cKO mice. Scale bars: 10 µm (A-D).

**Figure 4:**
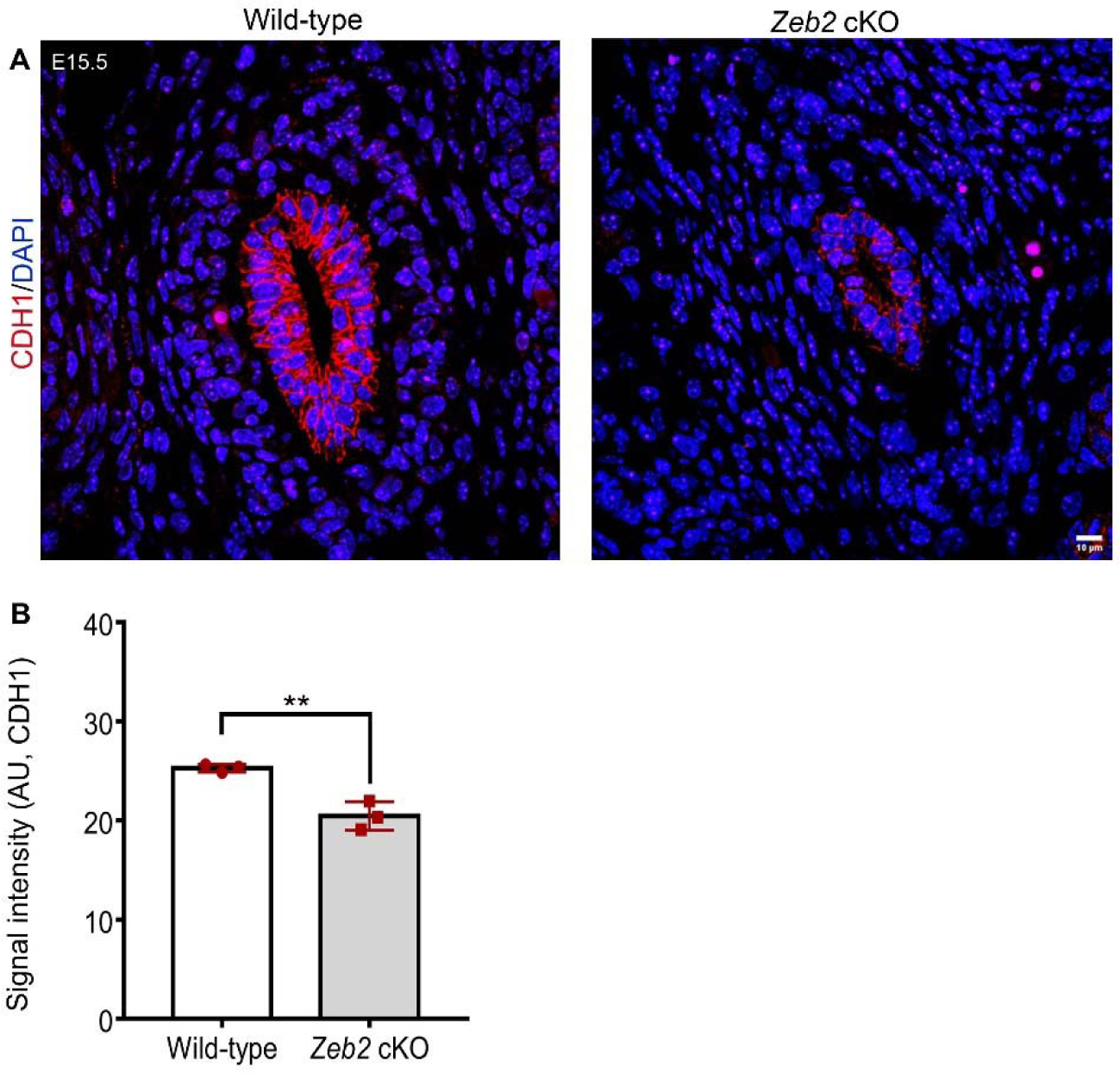
urothelium cell differentiation is altered in *Zeb2* cKO mice. (A) CDH1^+^ urothelium cells are underdeveloped in *Zeb2* cKO mice compared to wild-type littermate controls. (B) Quantification of CDH1 signal intensity in panel A. Scale bars: 10 µm (A).

### Loss of ZEB2 alters the expression of key transcription factors required for proper ureteral SMC differentiation and development

Previously, it has been shown that TBX18 (Gene ID: 9096) and SOX9 (Gene ID: 6662) are essential for ureteral SMC layer development, and loss of TBX18 or SOX9 leads to the absence of ureteral SMCs resulting in hydroureter [5, 31]. So, we performed immunostaining and examined TBX18 and SOX9 expressions to understand the molecular phenotypic changes in the *Zeb2* cKO ureters. We found a significant reduction in TBX18 expression in the *Zeb2* cKO mouse ureter at E14.5 and E15.5 (Figure 5A-B). Interestingly, SOX9 expression was significantly upregulated in the E15.5 *Zeb2* cKO mouse ureter as compared to their wild-type littermate controls (Figure 5C-D). These data suggest that loss of ZEB2 perturbs the expression of key transcription factors such as TBX18 and SOX9 which are critical for proper ureteral SMC differentiation and development during ureter formation. We also checked if the mesenchymal cell condensation was disturbed in *Zeb2* cKO mice as TBX18 is required for ureteral mesenchymal cell condensation [5]. Histological examination showed that ureteral mesenchymal cells were not condensed properly but instead loosely packed in *Zeb2* cKO pups as compared to the wild-type (Figure 6A-B).

**Figure 5.**
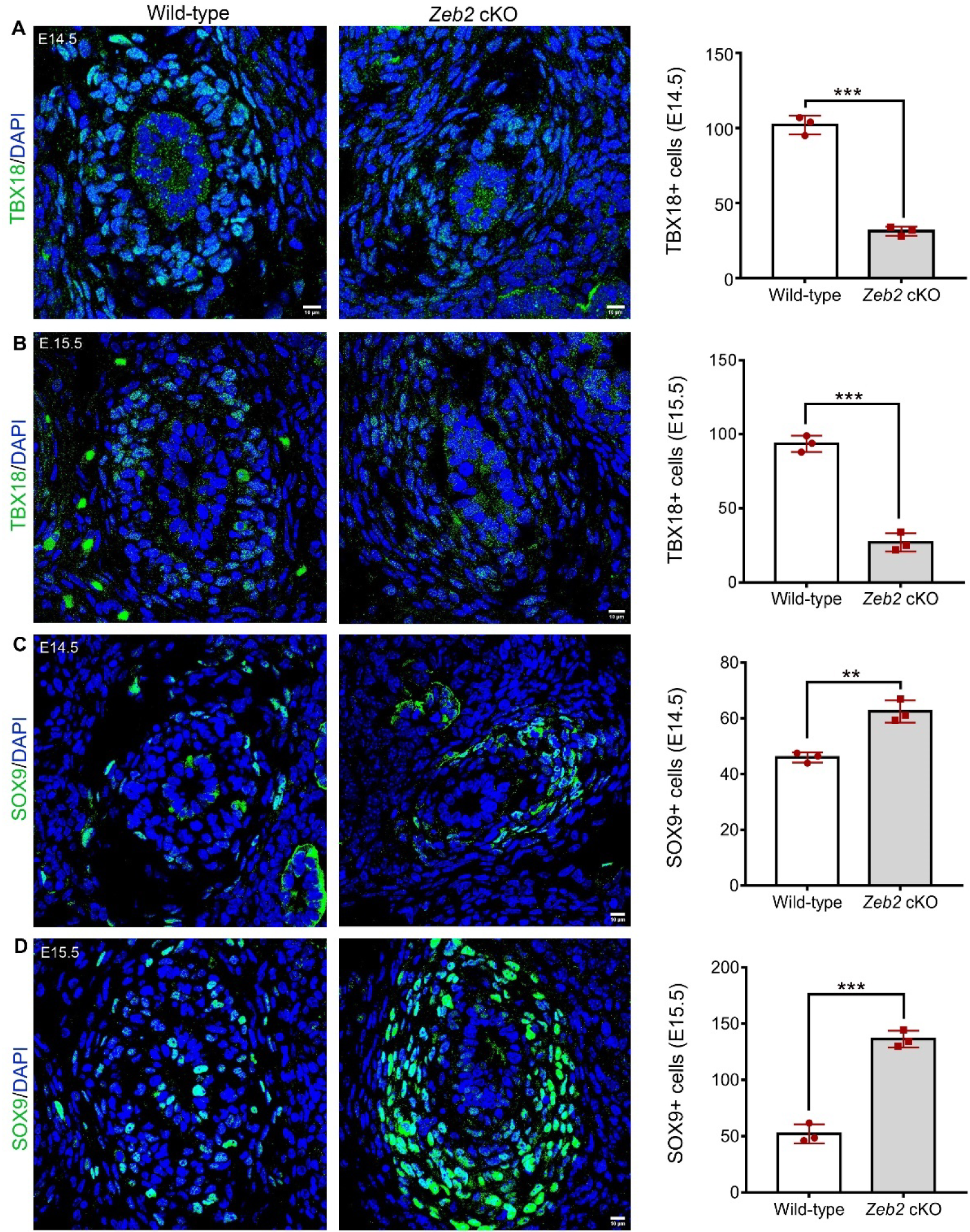
Loss of ZEB2 alters the expression of key factors required for proper ureteral SMC differentiation and development. (A-B) Immunostaining shows decreased TBX18 (green) expression in ureteral mesenchymal progenitors at E14.5 and E15.5 in *Zeb2* cKO mice compared to wild-type controls. (C-D) Immunostaining shows increased SOX9 (green) expression in ureteral mesenchymal progenitors at E14.5 and E15.5 in *Zeb2* cKO mice compared to wild-type controls. Scale bars: 10 µm (A-D).

**Figure 6:**
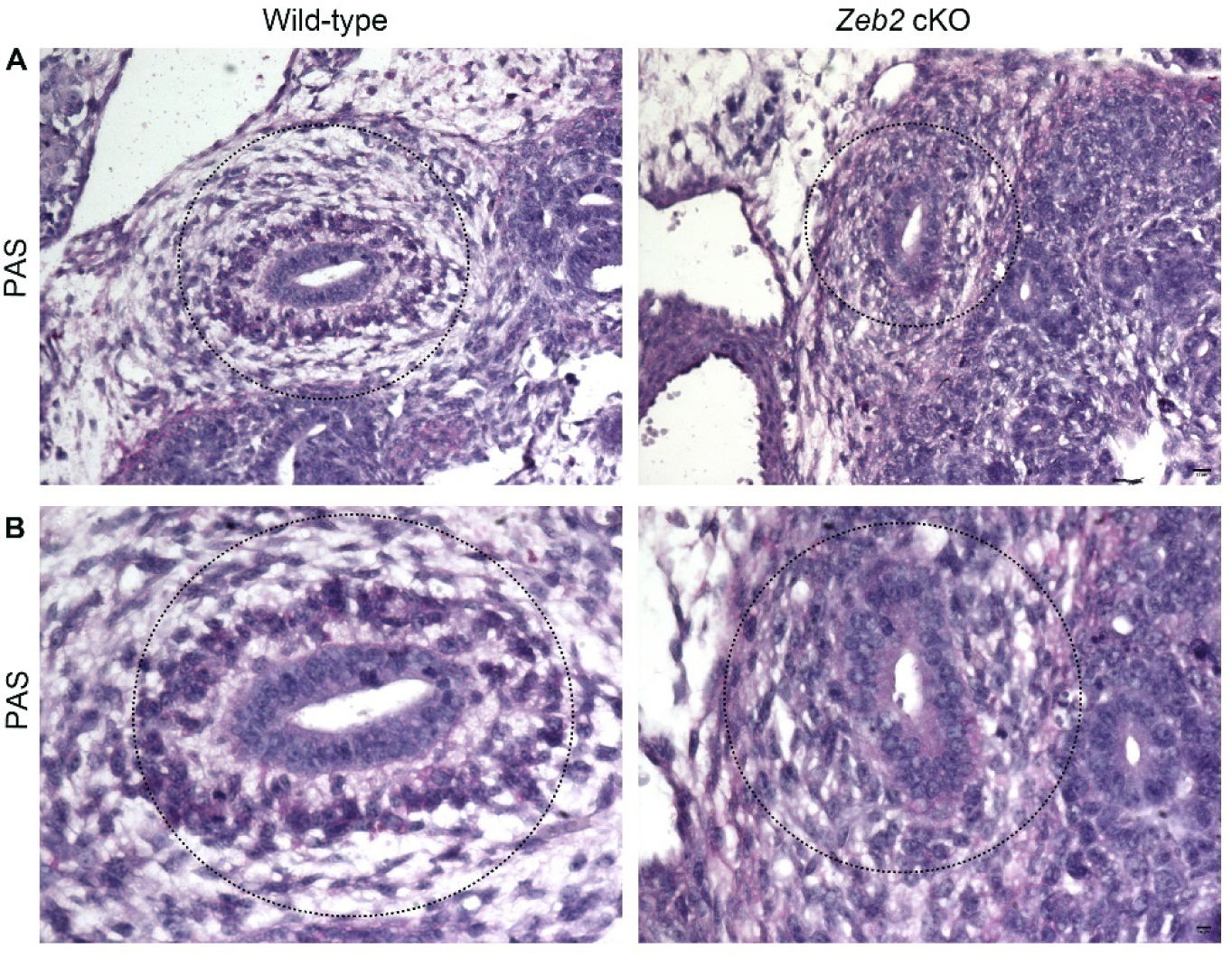
Ureteral mesenchymal cells are not appropriately condensed in the absence of ZEB2. (**A-B**) PAS staining shows normal condensation of ureteral mesenchymal cells in the wild-type mice at E14.5 but it is packed loosely in *Zeb2* cKO mice. Dotted lines encircle the ureter. Scale bars: 25 µm (A) and 10 µm (B).

## Discussion

Mowat-Wilson Syndrome (OMIM #235730; MWS) is caused by heterozygous mutations or deletions in the *ZEB2* gene, and more than half of MWS patients have urinary tract anomalies, including duplex kidney, pelvic kidney, VUR, and hydronephrosis [8–14]. In this report, we found that ZEB2 is expressed in the TBX18^+^ ureteral mesenchymal cells, and deletion of *Zeb2* by *Tbx18Cre*^+^ leads to hydroureter, hydronephrosis, and obstructive uropathy. We further found that *Zeb2* cKO mice do not develop the ureteral SMC layer but have an expanded layer of tunica adventitia cells. Mechanistically, we found that loss of ZEB2 leads to a reduced expression of transcription factor TBX18 but an increased expression of SOX9 in the ureteral mesenchymal cells, both are critical for ureteral SMC formation (Figure 7). These findings, hence, provide new molecular insights into the novel role of ZEB2 in the differentiation and development of ureteral SMCs.

**Figure 7:**
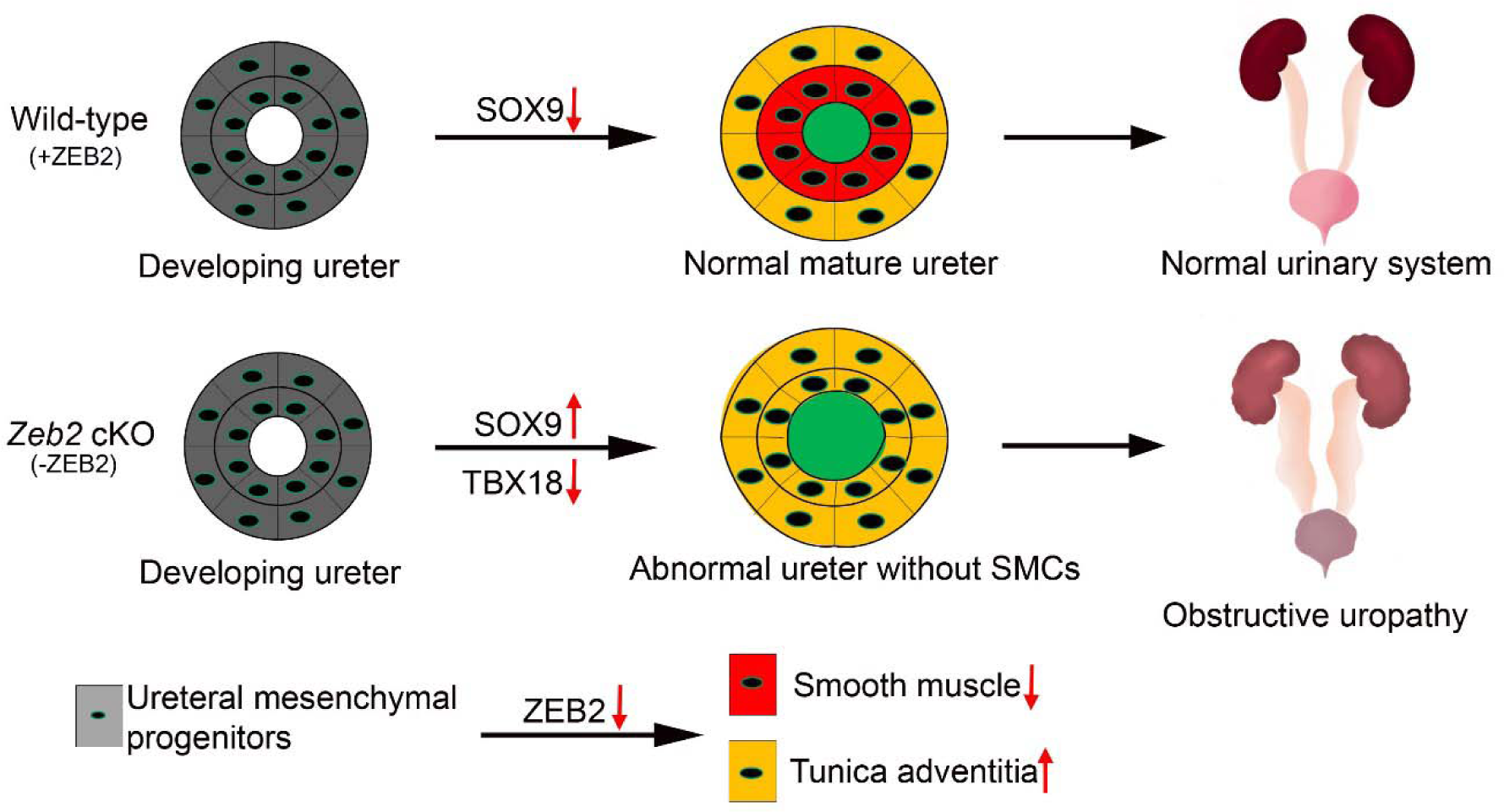
Proposed Model: ZEB2 signaling is essential for ureteral smooth muscle cell differentiation and maintenance

There is a paucity of information on molecular signals regulating ureteral SMC development [2, 3]. Only a few factors controlling ureter development have been identified so far, including Sonic Hedgehog signaling, WNT signaling, *Tbx18*, *Sox9*, *Bmp4*, *Six1*, and *Smad4* [1–3, 5, 28, 32–37]. *Tbx18* regulates the specification and development of the ureteric mesenchyme and *Tbx18^−/−^* knockout mice do not develop the SMC layer [5]. SOX9 is also crucial for ureteral mesenchymal cell differentiation, and loss of SOX9 also leads to the absence of SMCs [31]. Interestingly, *Zeb2* cKO mice also do not develop SMCs and have decreased expression of TBX18 but increased expression of SOX9 in the developing ureteral mesenchymal cells. Previously, it has been shown that ZEB2 antagonizes the inhibitory effect of SOX2, a SOX family protein, on Schwann cell differentiation and myelination [38, 39]. ZEB2 also controls SMC phenotypic transition through chromatin remodeling and loss of ZEB2 leads to differentially open chromatins for mature SMC regulatory markers such as *Cnn1*, *Myh11*, and *Myocd*, and chondrogenesis-related genes such as *Spp1*, *Sox9*, and *Col2a1* [40]. Our findings in this study discover ZEB2 as a novel ureter regulatory gene and suggest that ZEB2 might antagonize the inhibitory effect of SOX9 during ureteral SMC differentiation and development by altering chromatin accessibility. Further studies are needed to elucidate how ZEB2 interacts with TBX18 and SOX9 during ureter development and downstream genes that ZEB2 regulates in the developing ureter.

We also observed an expansion of the ureteral tunica adventitia cells in the *Zeb2* cKO mice in addition to SMC loss in the developing ureter. TBX18^+^ mesenchymal progenitors can subdivide into the inner and outer mesenchymal cells during ureter development. The inner mesenchymal cells further differentiate into the functional SMC layer to support ureter unidirectional peristalsis and prevent ureter dilation, whereas the outer compartment of TBX18^+^ mesenchymal cells gives rise to tunica adventitia cells [4]. Expansion of tunica adventitia cells in *Zeb2* cKO mice might be due to failure of the TBX18^+^ mesenchymal progenitors separation into inner and outer compartments in the absence of ZEB2. As a result, all TBX18^+^ mesenchymal progenitors differentiate into tunica adventitia cells. Alternatively, loss of the SMC layer in the *Zeb2* cKO may stimulate the expansion of the ureteral tunica adventitia layer. Thirdly, this unexpected cellular phenotype of expanded ureteral tunica adventitia layer could also be driven by upregulated SOX9 expression in the ureteral mesenchymal cells as prolonged expression of SOX9 in the ureteral mesenchyme reduces smooth muscle gene expression and increases the expression of extracellular matrix (ECM) components as previously reported [31]. Lastly, a recent study also suggests that SMC-specific loss of ZEB2 results in an inability to turn off contractile programming and transitioning SMCs to take on a fibroblast-like phenotype [40]. Further studies are needed to confirm if ZEB2 specifically regulates early ureteral mesenchymal cell specification during ureter development and if expanded ureteral tunica adventitia layer phenotype is merely a compensatory response to SMC loss in *Zeb2* cKO mice.

In past years, much progress has been made in understanding the role of ZEB2 during nervous system development [18, 41]. However, the role of ZEB2 in the urinary system is poorly understood, even though more than 50% of MWS patients have been reported as having urinary tract anomalies. Our current work not only demonstrates the important role of ZEB2 in ureter development but also provides molecular genetic insights into some aspects of MWS-associated phenotypes in the urinary tract.

In summary, we found that loss of ZEB2 in ureteral mesenchymal cells leads to ureteral SMC loss, which eventually causes hydroureter, hydronephrosis, and obstructive uropathy (Figure 7). Our data suggest that ZEB2 is essential for ureteral SMC differentiation and development during ureter development. Our study also indicates that urinary tract anomalies are the primary phenotype in MWS patients and that MWS patients should be evaluated for urinary tract disorders such as hydroureter, hydronephrosis, and obstructive uropathy.

## Materials and Methods

### Animals

All animal experiments were performed according to the guidelines issued by the Institutional Animal Care and Use Committee of Boston University Medical Campus at Boston University (approved IACUC protocol number: PROTO201800056_TR01). *Zeb2* floxed mice [26] and *Tbx18Cre^+^* mice [27] were genotyped using primers previously reported. ZEB2-EGFP knock-in reporter mouse was studied previously [25]. Eight-week-old *Zeb2*^flox/+^;*Tbx18Cre^+^* mice were bred with *Zeb2*^flox/flox^ mice to generate *Zeb2*^flox/flox^;*Tbx18Cre^+^* homozygous conditional knockout mice. All mice had free access to drinking water and a standard rodent diet ad libitum. Animals of both sexes in mixed C57BL/6 x 129 genetic backgrounds were used in this study. Wild-type littermates were used as controls. Timed-pregnant females were confirmed by semen plug and designated at embryonic day 0.5 (E0.5) of gestation and were separated from the breeding cage.

### Tissue Preparation and Histology

Gross urinary tracts of newborn *Zeb2* cKO mice and wild-type littermates were dissected and imaged with an Olympus inverted dissecting microscope (Olympus, USA). For histological analysis, urinary tracts were fixed in 4% paraformaldehyde or 10% neutral buffered formalin at 4°C overnight and processed for paraffin embedding following standard protocols. Serial kidney sections were cut and stained using a Periodic acid Schiff (PAS) stain kit (# 24200, Polysciences, Inc. USA), according to the manufacturer’s instructions. Slides were examined with an Olympus upright light microscope and photographed using an Olympus DP72 digital camera.

### Immunofluorescence Staining and Confocal Microscopy

Embryos were harvested from timed-pregnant females at embryonic day (E) 13.5, 14.5, 15.5, and 16.5. Whole embryos were fixed in 4% formaldehyde overnight at 4°C followed by incubation in 30% sucrose overnight at 4°C, embedded in optimal cutting temperature (OCT) compound (Tissue-Tek, Sakura Finetek), and cryosectioned at 8-10 μm. Frozen sections were permeabilized with PBS containing 0.1% Triton X-100 for 10 minutes and blocked in 5% serum-blocking buffer or background buster (NB306, Innovax biosciences, USA) for 1 hour at room temperature. Primary antibodies were incubated at 4°C overnight, followed by secondary antibodies at room temperature for 1 hour. Following primary antibodies were used: ZEB2 (1:100, HPA003456; Sigma-Aldrich, USA or SC-271984, Santa Cruz Biotechnology, USA), TBX18 (1:100, ab115262, Abcam, USA), E-cadherin (1:50, Clone 36/610181; BD Biosciences, USA), ACTA2 (1:200, A2547; Sigma-Aldrich, USA), Cytokeratin (1:100, C2562, Sigma-Aldrich, USA), TAGLN (1:200, ab14106, Abcam, USA), FOXD1 (1:100 dilution, sc-47585; Santa Cruz Biotechnology USA), POSTN (1:100, ab14041, Abcam, USA), SOX9 (1:200, AB5535, Millipore, USA), GFP (GFP-1020, 1:100, Aves Labs, USA or A11122, 1:250; Invitrogen, Carlsbad, CA). Secondary antibodies were goat anti-mouse IgG2a AF594, goat anti-mouse IgG1 AF488, goat anti-rabbit IgG AF488, goat anti-rabbit IgG AF494, and donkey anti-rabbit IgG AF594. Sections were stained with DAPI (4’,6’-’diamidino-2-phenylindole) and mounted in Vectashield antifade mounting medium (#H-1000; Vector Labs, USA). Images were captured by Zeiss LSM 700 confocal microscope (Zeiss, Germany) and analyzed with ImageJ software (National Institutes of Health, USA).

### Image quantification

Quantification of images was performed using ImageJ on random images (*n* = 6 per group). Cell counter or particle analysis plug-in tools were used to quantify the cell number. For intensity quantification, the percentage of total pixel area was measured.

### Statistics

All statistical analyses were performed using GraphPad Prism statistical software (version 7.0, GraphPad, San Diego, CA) with *P* < 0.05 as statistical significance. Wild-type and Zeb2 cKO mice group data were compared using a two-tailed unpaired Student’s *t-test*. Data are represented as mean ± SEM. A minimum of three mice were used for each analysis group unless stated otherwise.

## Acknowledgments

We thank Drs. David J. Salant, Laurence Beck, and Hila Milo Rasouly for their helpful discussion, as well as comments from colleagues of the Boston University Academic Writing program and the Kidney Research Program of the Nephrology Section. We thank Boston University’s analytical instrumentation core, cellular imaging core, and confocal microscopy facility for sample and tissue analysis. This work is supported by research grants from NIH/NIDDK grants R01DK078226, R01DK133940 (WL), a DOD grant E01 HT9425-23-1-1058 (WL), and a BU CTSI Integrated Pilot Grant funded by the Department of Medicine (SK, WL). This work is also supported by the Career Development Award from the American Heart Association (SK, https://doi.org/10.58275/AHA.24CDA1272560.pc.gr.193662), The Department of Medicine Career Investment Award (SK), and the Boston University Undergraduate Research Opportunities Program (HP, EJL, JJ, EZ, YJ). This publication is also supported in part by the National Center for Advancing Translational Sciences, National Institutes of Health, through BU-CTSI Grant Number 1UL1TR001430. Its contents are solely the authors’ responsibility and do not necessarily represent the official views of the NIH.

## Author Contributions

SK and WL conceived and designed the experiments. SK, HP, KY, EJL, JJ, EZ, PS, YJ, and YN performed the experiments and acquired the data. SK, XF, YN, YH, and WL analyzed the data. WL and YH provided reagents/materials/analysis tools. SK and WL wrote the manuscript. All authors reviewed and approved the final manuscript.

## Data Sharing Statement

All authors agree to share data.

